# Microstructures of Ant Chemosensory Sensilla Support a Dual Function in Detecting Both Volatile and Contact-Mediated Cues

**DOI:** 10.1101/2022.06.05.494864

**Authors:** Hannah R. Gellert, Daphné C. Halley, Zackary J. Sieb, Jody C. Smith, Gregory M. Pask

## Abstract

Ants and other eusocial insects emit and receive chemical signals to communicate important information within the colony. In ants, nestmate recognition, task allocation, and reproductive distribution of labor are largely mediated through the detection of cuticular hydrocarbons (CHCs) that cover the exoskeleton. With their large size and limited volatility, these CHCs are believed to be primarily detected through direct contact with the antennae during behavioral interactions. Here we use scanning electron microscopy to investigate the unique morphological features of CHC-sensitive basiconic sensilla of two ant species, the black carpenter ant *Camponotus pennsylvanicus* and the Indian jumping ant *Harpegnathos saltator*. These basiconic sensilla possess an abundance of small pores typical of most insect olfactory sensilla, but also have a large concave depression at the terminal end. Basiconic sensilla are enriched at the distal segments of the antennae in both species, further supporting their proposed role in contact chemosensation. A survey of these sensilla across other ant subfamilies shows varied microstructures at their tips, but each possess surface textures that would also increase sensory surface area. These unique ant chemosensory sensilla represent yet another example of how specialized structures have evolved to serve the functional requirements of eusocial communication.

## Introduction

Reliable communication among individuals is of the utmost importance in successful animal societies. In eusocial insects, this communication can employ several sensory modalities and drive a wide range of colony behaviors, as well as maintain the division of labor among different castes. The popular waggle dance of honeybees recruits foraging workers within the hive using auditory, vibrational, chemical, and tactile signals^1^. Ants rely heavily on chemical communication to signal foraging trails, detect invading non-nestmates, and maintain the reproductive hierarchy^2^. Chemical or genetic manipulation of these chemosensory communication systems can trigger nestmate aggression, disrupt colony social behaviors, and even decrease reproductive success^3–6^.

Cuticular hydrocarbons (CHCs) provide a desiccation-resistant barrier in terrestrial insects but have been co-opted by eusocial insects to serve as social cues^7,8^. A structurally diverse range of CHCs in ants, bees, wasps, and termites have been implicated in nestmate recognition, reproductive division of labor, and task distribution^9– 16^. For example, a queen ant has a specific CHC profile that reflects her colony identity and reproductive status, with a matching CHC profile on her eggs that can distinguish them from those of workers^17^. The majority of CHCs identified on ants tend to have chain lengths of 25-35 carbons with varying levels of unsaturation and methyl branching^18^. At these sizes, ant CHCs are believed to have substantially low volatility and likely function in close-or near-contact interactions^19^.

Antennal detection of CHCs is mediated by specialized sensory hairs, or sensilla, that has been confirmed by electrophysiology in several ant species^11,20–22^. Specifically, large basiconic sensilla are sensitive to a wide range of CHCs and house greater than 100 olfactory receptor neurons (ORNs)^23^. The genes expressed in these ORNs are likely from the odorant receptor (OR) family, as heterologous expression studies have characterized ant ORs as highly specific CHC detectors^24,25^. In these functional studies, the volatility of CHCs is augmented by either gas chromatography or direct application of heat^21,22,24,25^. However, it is still unclear if CHC profiles are reliably detected by the antennae as volatile cues or if physical contact is necessary.

Here, we use scanning electron microscopy (SEM) to provide insight into how morphology may facilitate antennal discrimination of CHC cues. We examined two distantly related ant species: the black carpenter ant, *Camponotus pennsylvanicus*, where colonies consist of morphologically distinct castes and an established reproductive, and the Indian jumping ant, *Harpegnathos saltator*, that has a single morphological worker caste and displays reproductive plasticity. We show that basiconic sensilla are multiporous, characterized by several smaller pores and a larger concave depression at the tip. These sensilla are predominantly enriched in the distal ends of the antennae, the regions that contact other ants in observed social interactions. In a survey of other major ant subfamilies, similar basiconic sensilla possessing concave microstructures are found in only some species, with others appearing to have other adaptions that could also increase sensory surface area. Our findings suggest that the evolution of ant chemosensory appendages has resulted in structural adaptations in sensilla that serve a dual function for both contact-mediated recognition of social cues and airborne detection of volatile odorants.

## Results and Discussion

SEM analysis revealed that the basiconic sensilla of *C. pennsylvanicus* feature a large concave depression surrounded by several smaller pores at the distal end of the sensillum (Figure 1a, c, e, g). In *C. pennsylvanicus* minor workers, the small multiporous openings have a mean diameter of 0.07µm ± 0.009 µm; meanwhile, the terminal depression is approximately 1.4µm ± 0.11µm in width and 0.39µm ± 0.014µm in height. This sensillar microstructure was conserved in various female morphs of *C. pennsylvanicus*: nanitics (smaller first workers), minors, intermediates, majors, and queens (data not shown). Furthermore, basiconic sensilla on the antennae of the ponerine ant, *H. saltator*, also possessed a concave depression at the tip (Figure 1b, d, f, h). In *H. saltator* workers, the small multiporous openings are 0.09µm ± 0.013µm in diameter; meanwhile, the terminal depression is approximately 1.4µm ± 0.11µm in width and 0.39µm ± 0.06µm in height.

**Figure 1.**
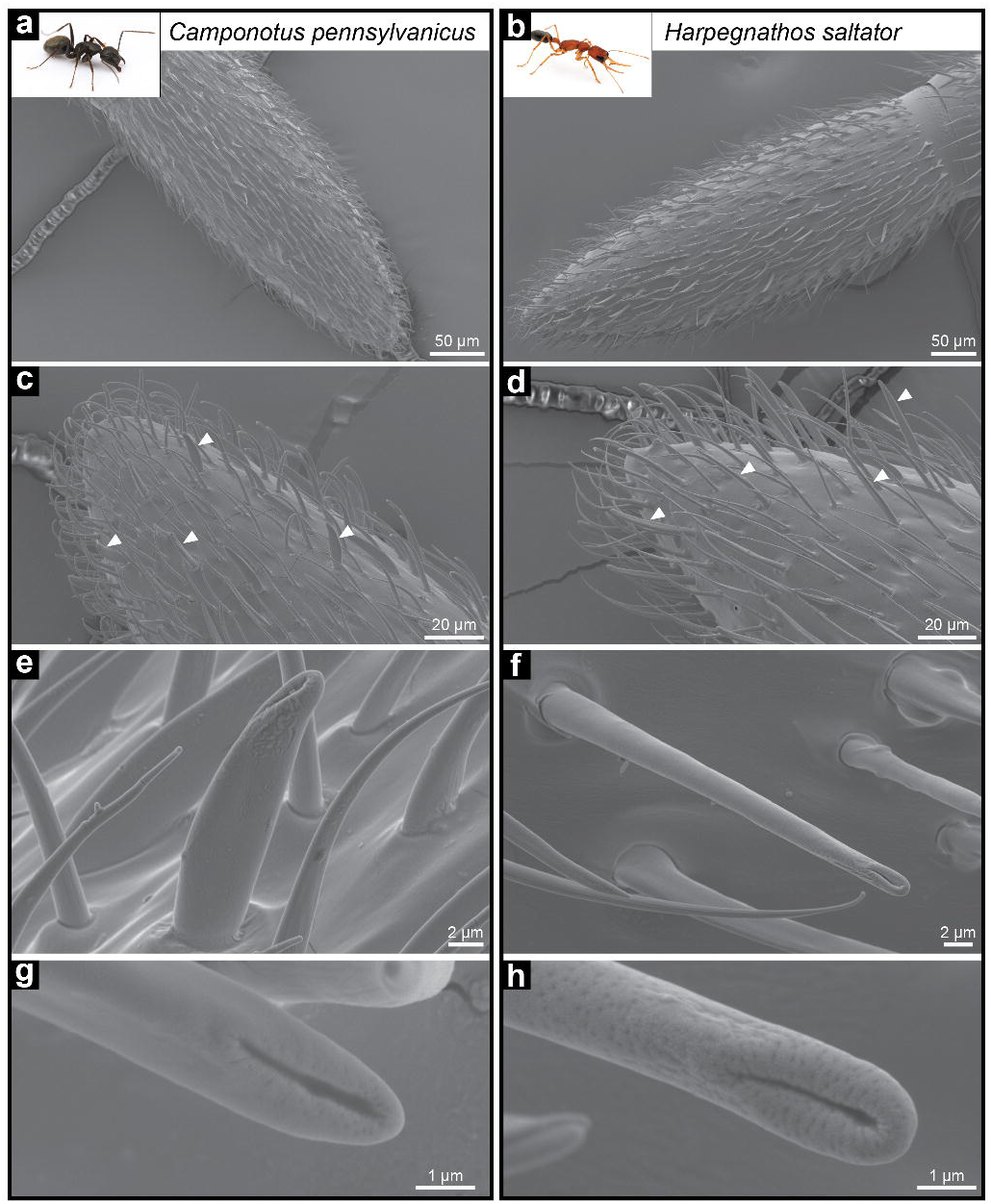
Morphology of chemosensory sensillum in *C. pennsylvanicus* and *H. saltator*. Representative images of a *Camponotus pennsylvanicus* minor worker (a, c, e, g) and *Harpegnathos saltator* worker (b, d, f, h). Panels depict the full 13th antennal segment (a, b) and its distal tip with some basiconic sensilla highlighted by arrowheads (c, d). Images of a basiconic single sensillum (e, f) and the terminal end (g, h) depict a large concave depression and several smaller pores. Inset ant images are provided courtesy of Alex Wild (alexanderwild.com).

From SEM imaging it is unclear whether the visible depression at the end of the basiconic sensilla in *C. pennsylvanicus* and *H. saltator* is merely a concave tip or a terminal pore that connects directly to the sensillar lymph. A concave depression would increase surface area for more efficient contact with surfaces (nestmate cuticle or others) during antennation. A terminal pore would be similar to uniporous gustatory sensilla found in insect taste appendages, where tastants enter the lymph through contact with a single large opening at the tip of the sensillum^26^. CHCs contacted by basiconic sensilla during antennation could traverse this terminal pore, enter the sensillum lymph where it is solubilized by a family of odorant-binding and chemosensory proteins, and interact with the olfactory receptor neurons (ORNs)^27^. Further imaging with transmission electron microscopy could investigate this question of basiconic sensillum tip ultrastructure and potential lymph continuity.

Our structural observations of ant basiconic sensilla seem to align with previous functional and neuroanatomical studies that have characterized these sensilla as detectors of CHCs and general odorants^11,20–23^. Electrophysiological recordings from several ant species have found that ORNs within basiconic sensilla respond to a wide range of linear and methyl-branched hydrocarbons, including many that are found in ant cuticular extracts. Basiconic ORNs are also sensitive to general odorants, such as alcohols, esters, and acids, that are relatively smaller and more volatile than CHCs^21,22^. It is worth noting that ant basiconic sensilla are innervated by ∼100 ORNs, whereas other insects typically have 1-4 ORNs per sensillum^23,28–31^. A sensillum housing so many ORNs that respond to diverse chemical stimuli perhaps benefits from specialized structural features that can enable the efficient detection of both volatile and contact-mediated cues.

We then imaged full-length antennae of both *C. pennsylvanicus* and *H. saltator* to quantify the abundance of basiconic sensilla on each segment (Figure 2). In all five female *C. pennsylvanicus* castes and *H. saltator* workers, basiconic sensilla were significantly enriched in the distal segments and accounted for >68% of total variation (Figure 2, Table 1, two-way ANOVA, p<0.0001 for all datasets). Interestingly, a significant bias toward ventral position of these sensilla was found in *C. pennsylvanicus* minor and major workers, but no other samples (Supplementary Figure S1, two-way ANOVA, p=0.0034 for minors and p<0.0001 for majors). At the extremes of the funiculus, the 13th segments of *C. pennsylvanicus* minors and *H. saltator* workers had 47.3 ± 4.8 and 63.8 ± 3.7 basiconic sensilla, respectively, and decreased proximally along the antenna until the 3rd segment, where no basiconic sensilla were present across both species and all castes of *C. pennsylvanicus*. Increased basiconic sensilla presence at the distal end and/or ventral surface of an antennae coincides with the regions that predominantly contact other ants during typical social interactions, providing further support for a model of contact-or near contact-mediated recognition of CHCs^32^.

**Figure 2.**
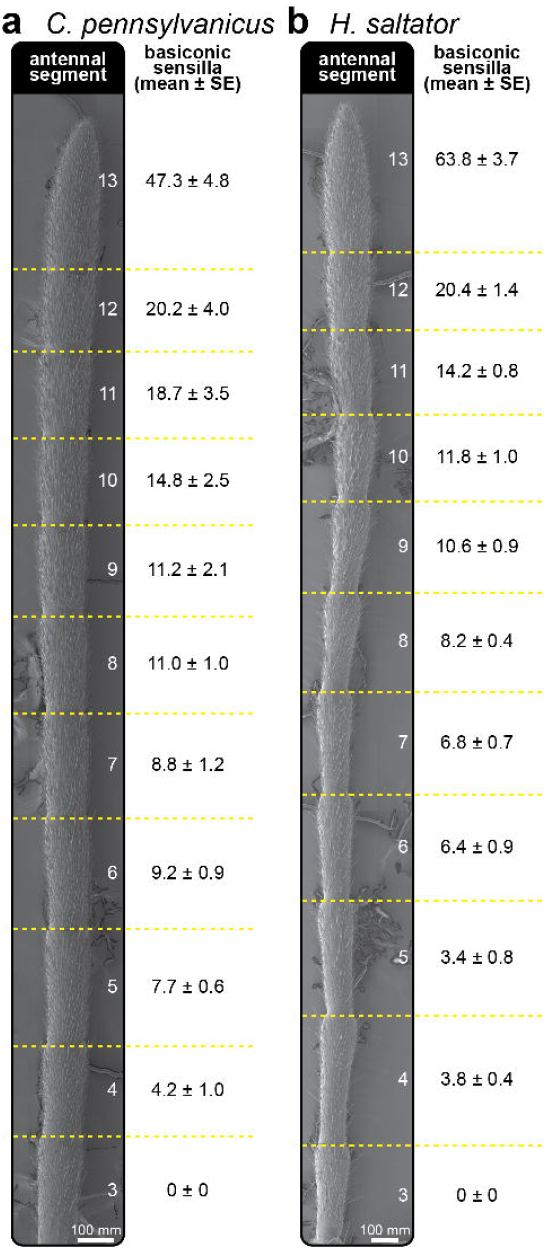
Distal abundance of basiconic sensilla across two distantly related ant species. Antennal images (left) and basiconic sensilla counts (right) of a *C. pennsylvanicus* minor worker (a) and *H. saltator* worker (b). Sensillum counts represent the average and standard error of the dorsal and ventral total per individual (*n* = 5-6)

**Table 1.** Antennal distribution of basiconic sensilla across different *C. pennsylvanicus* castes. Sensillum counts represent the average and standard error of the dorsal and ventral total for each antennal segment (*n* = 5-6).

| **Caste** | **Antennal Segment** |  |  |  |  |  |  |  |  |  |  |
| --- | --- | --- | --- | --- | --- | --- | --- | --- | --- | --- | --- |
|  | 3 | 4 | 5 | 6 | 7 | 8 | 9 | 10 | 11 | 12 | 13 |
| *nanitic* | 0.0 ±<br>0.0 | 3.3 ±<br>0.6 | 3.8 ±<br>1.2 | 5.5 ±<br>1.3 | 6.2 ±<br>1.0 | 7.5 ±<br>1.1 | 8.0 ±<br>1.7 | 9.5 ±<br>2.6 | 10.2 ±<br>2.8 | 11.0 ±<br>3.5 | 37.0 ±<br>6.2 |
| *minor* | 0.0 ±<br>0.0 | 4.2 ±<br>1.0 | 7.7 ±<br>0.6 | 9.2 ±<br>0.9 | 8.8 ±<br>1.2 | 11.0 ±<br>1.0 | 11.2 ±<br>2.1 | 14.8 ±<br>2.5 | 18.7 ±<br>3.5 | 20.2 ±<br>4.0 | 47.3 ±<br>4.8 |
| *intermediate* | 0.0 ±<br>0.0 | 4.2 ±<br>0.4 | 6.6 ±<br>0.4 | 8.0 ±<br>0.7 | 9.0 ±<br>1.2 | 10.0 ±<br>1.0 | 12.0 ±<br>0.5 | 14.8 ±<br>1.2 | 17.6 ±<br>1.2 | 17.6 ±<br>1.2 | 50.0 ±<br>2.9 |
| *major* | 0.0 ±<br>0.0 | 3.8 ±<br>0.9 | 6.6 ±<br>0.9 | 9.6 ±<br>0.5 | 9.8 ±<br>0.4 | 13.2 ±<br>0.9 | 14.2 ±<br>1.7 | 19.4 ±<br>2.4 | 21.6 ±<br>2.6 | 24.8 ±<br>4.6 | 61.2 ±<br>7.6 |
| *queen* | 0.0 ±<br>0.0 | 2.4 ±<br>0.4 | 6.4 ±<br>0.5 | 8.0 ±<br>1.1 | 9.0 ±<br>1.4 | 11.8 ±<br>0.7 | 14.8 ±<br>0.7 | 16.2 ±<br>1.4 | 22.4 ±<br>1.2 | 25.0 ±<br>1.6 | 64.8 ±<br>2.6 |

\*denotes subfamilies examined in this study

We expanded our characterization of these unique basiconic sensillar microstructures to other subfamilies of ants (Figure 3). Notably, distally enriched basiconic sensilla in *Formica exsectoides* (subfamily Formicinae) and *Pogonomyrmex occidentalis* (subfamily Myrmicinae) possessed similar concave features at their ends along with numerous smaller pores (Figures 4a, b). However, in *Atta cephalotes* (subfamily Myrmicinae), *Linepithema humile* (subfamily Dolichoderinae), *Tapinoma sessile* (subfamily Dolichoderinae), and *O. biroi* (subfamily Dorylinae) no large depression was observed at the tip of basiconic sensilla (Figures 4c-f). Instead, the sensilla displayed rounded and multiporous ends with some possessing elaborate grooves along the distal surface. Grooves on the surface of olfactory sensilla have been observed in several insects and have been theorized to aid in directing hydrophobic odorants into pores^33,34^. It is possible that the sensillar grooves represent another morphological strategy to increase sensory surface area and retain CHCs. Although there is no clear evolutionary pattern on the presence of terminal depressions of basiconic sensilla, further imaging across the Formicidae subfamilies may reveal a social, chemical, or morphological role.

**Figure 3.**
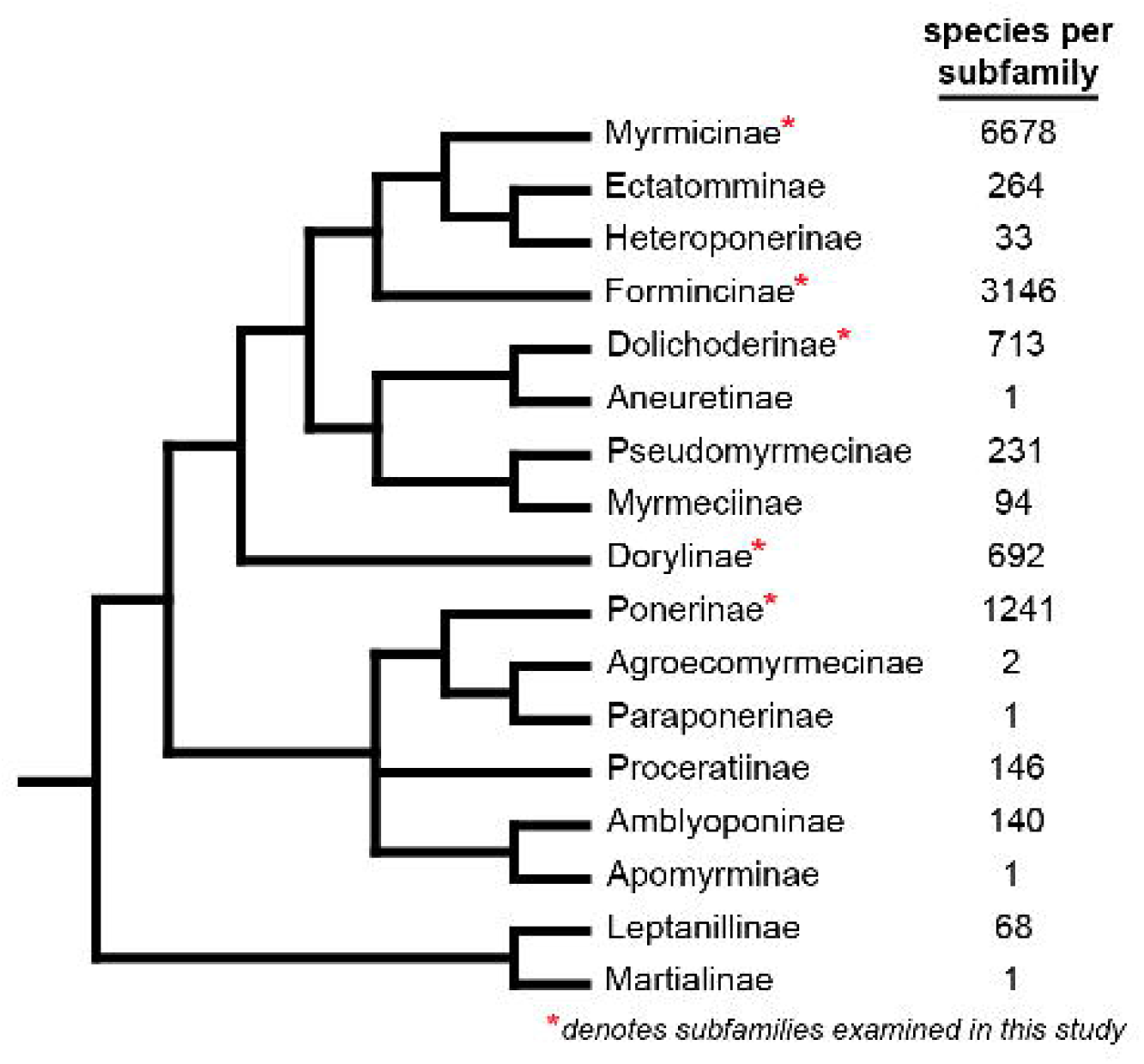
Phylogenetic relationships between ant subfamilies used in this study. Cladogram of ant subfamilies and species numbers created from data in Borowiec et al^42^.

**Figure 4.**
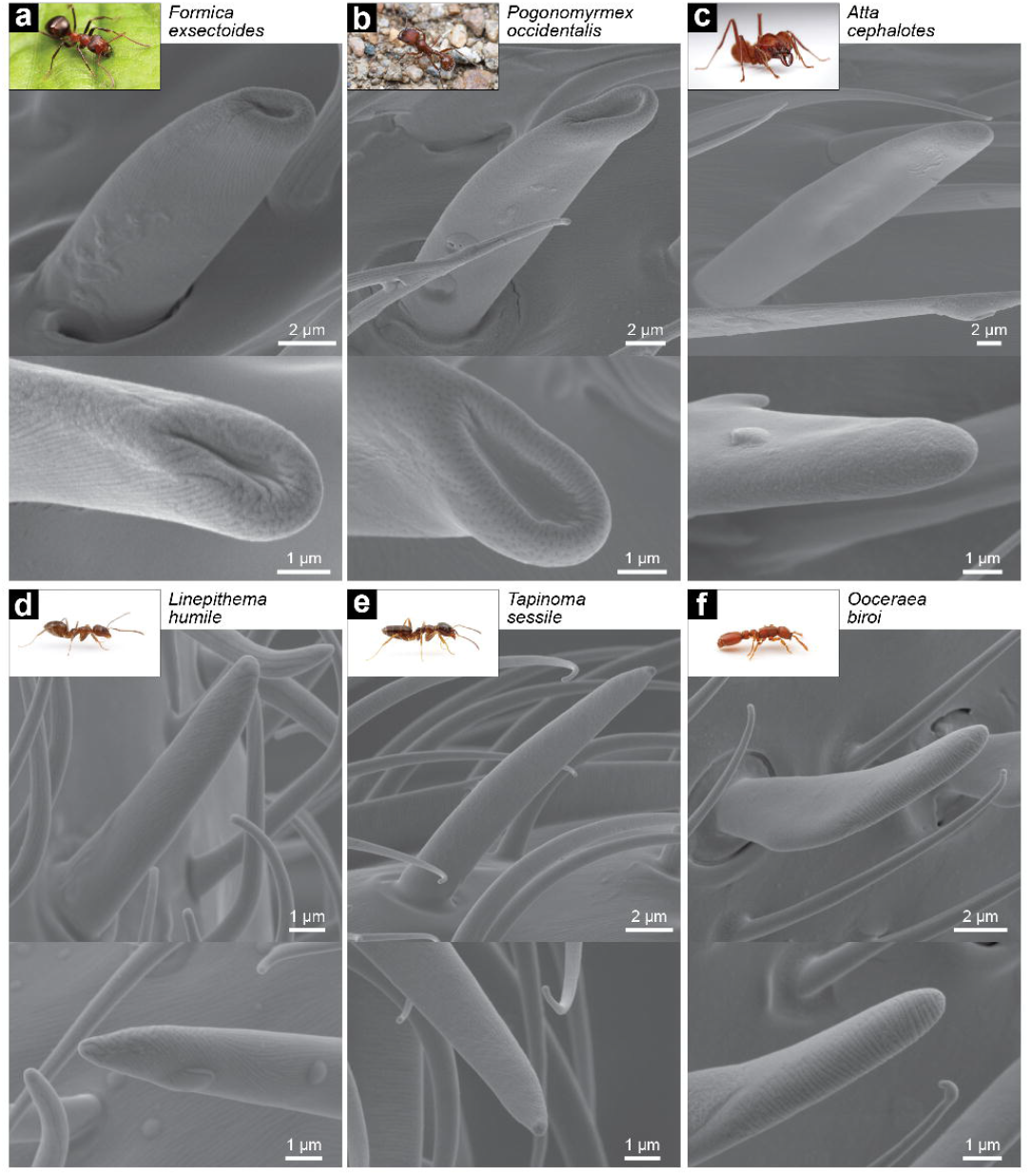
Basiconic sensillum morphology across major subfamilies of ants. Images of both the basiconic sensillum (top) and its tip (bottom) of (a) *Formica exsectoides* worker (subfamily Formicinae), (b) *Pogonomyrmex occidentalis* worker (subfamily Myrmicinae), (c) *Atta cephalotes* super major (subfamily Myrmicinae), (d) *Linepithema humile* worker (subfamily Dolichoderinae), (e) *Tapinoma sessile* worker (subfamily Dolichoderinae), and (f) Ooceraea biroi (subfamily Dorylinae). Inset ant images are provided courtesy of Alex Wild (alexanderwild.com).

Ant antennae across many species have been imaged by SEM before, yet these specific features of basiconic sensilla have not been previously described ^22,23,32,35,36^. Examination of previous images shows faint indications of the terminal concave depression, but higher accelerating voltages of 15-20kV result in deeper tissue penetration by the electrons, thus potentially causing a loss of surficial features^22,23^. In our case, reduction of the accelerating voltage to 5kV revealed clear depressions and provided increased resolution of the smaller olfactory pores. As advances in electron optics continue, lower accelerating voltages may increase the surficial resolving power of insect SEM imaging.

As a major sensory interface for insects, the varied structural aspects of cuticular protuberances serve specific roles in sensory function. For example, the long thin hairs of plumose mosquito antennae are firmly coupled to the antennal shaft and can efficiently transmit the wingbeat frequencies of conspecifics through the shaft to the Johnston’s organ^37^. Our morphological characterization of a specialized chemosensory sensillum in ants identifies features that align with known functional and behavioral aspects of CHC detection and nestmate recognition, with both the increased surface area of the terminal microstructure and the distal abundance of ant basiconic sensilla supporting contact recognition of CHCs. Although it has be shown close-range recognition of non-nestmate CHCs can occur without antennal contact^19^, recognition behaviors of free-moving ants begin with investigative antennation with repeated contact before a decision is made regarding acceptance or aggression^38–40^. Our findings support the further investigation of questions in non-model insects, as the identification of unique morphological structures can inform and support our understanding of well-documented behaviors.

## Methods

### Animals

All female castes of *Camponotus pennsylvanicus* and workers of *Formica exsectoides* were collected locally in Vermont. *Harpegnathos saltator* were obtained from an in-house laboratory colony. *Pogonomyrmex occidentalis* workers were purchased from TruBlu Supply. Workers from the following species were generously donated by the following individuals: *Atta cephalotes* (M. Gilbert, University of Pennsylvania), *Linepithema humile* (L. Martins and N. Tsutsui, University of California, Berkeley) *Tapinoma sessile* (G. Buczkowski, Purdue University), and *Ooceraea biroi* (W. Trible, Harvard University). Only female ants were imaged as male ants have already been shown to not possess basiconic sensilla^23,41^.

### Scanning Electron Microscopy (SEM)

#### Sample Preparation

For each insect anesthetized by CO_2_, the head was removed with antennae intact. The head and both antennae were sequentially washed in a watch glass with pure hexanes, pure acetone, and 95% ethanol, with each solvent applied only after the previous had fully evaporated. The left antenna was mounted on a ½” slotted head aluminum specimen mount, dorsal side up, then removed from the head. On the same mount, the right antenna was positioned ventral side up, then removed from the head. Samples were coated in gold-palladium using an Ernest Fullam Inc.EffaCoater Au-Pd Sputter Coater.

#### Imaging and Analysis

Images were acquired using a Tescan Vega 3 LMU Scanning Electron Microscope with an accelerating voltage of 5kV and beam intensity of 6. High-quality imaging of whole antennae was performed with automated scanning and post-hoc montage stitching using Tescan VegaTC software image snapper wizard.

Images were analyzed for the presence of basiconic sensilla throughout the antenna. For *C. pennsylvanicus* and *H. saltator*, basiconic sensilla are only present on the funiculus (segments 3-13); therefore, imaging and counting excluded the scape. Each segment of the antenna was numbered, with segment three being the proximal segment and segment thirteen being the distal segment at the tip of the antenna. Basiconic sensilla were counted only if the base of the sensillum was visible to prevent possible double counting of sensilla from the opposite side. This process was repeated for the dorsal and ventral sides, and then combined for an estimate of the total basiconic sensilla present on a single antenna.

Statistical analysis of basiconic sensilla counts was performed using Prism 9 (Graphpad). Two-way ANOVA tests were used to determine the effects of antennal segment and dorsal/ventral surface on sensillum abundance, with Bonferroni’s multiple comparisons tests for post-hoc comparisons between dorsal/ventral abundance at each segment.

## Acknowledgments

We would like to thank Michael Gilbert, Lourenço Martins, Neil Tsutsui, Grzegorz Buczkowski, W. Trible, and Daniel Kronauer for providing ant specimens and Alex Wild for sharing all ant images inset in Figures 1 and 3. We also thank Stephen Ferguson, Sarah Lower, Jason Pitts, and Hua Yan for insightful comments on the manuscript. We also thank the George I. Allen Trust for funding the acquisition of the Tescan Vega 3 LMU Scanning Electron Microscope and the Middlebury Undergraduate Research Office and Biology Department for providing student research support.

## Author Contributions

HRG, DCH, and GMP designed the experiments. HRG, DCH, and ZJS prepared and imaged antennal samples. HRG, DCH, ZJS, and JCS optimized imaging conditions. HRG and GMP performed data analyses and prepared figures and tables. HRG and GMP wrote the manuscript and all authors reviewed the manuscript.

## Data Availability Statement

Imaging data and analyses are available from the corresponding author upon request.

