## Supplementary Figure 1 for "Microstructures of Ant Chemosensory Sensilla Support a Dual Function in Detecting Both Volatile and Contact-Mediated Cues"

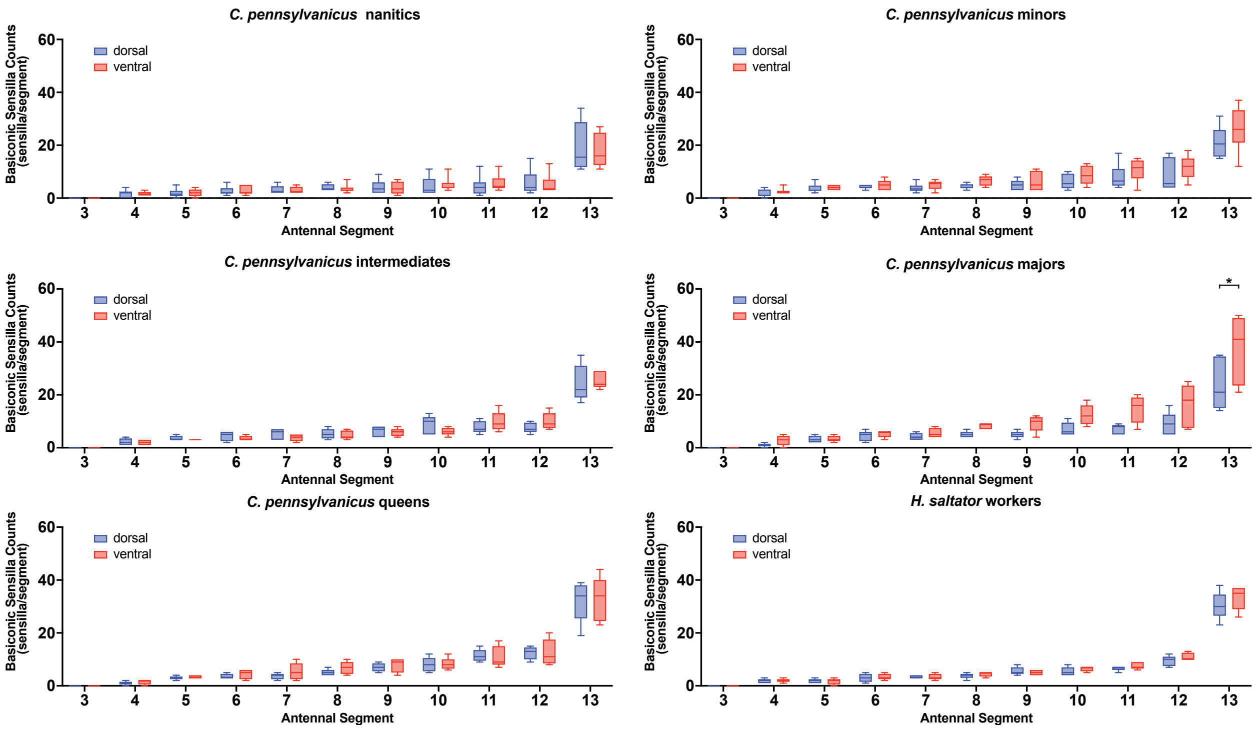


**Supplementary Figure S1. Abundance of Basiconic Sensilla by Antennal Segment and Dorsal/Ventral Surface.** In all datasets, the antennal segment had a significant effect on basiconic sensillum abundance (two-way ANOVA, $p<0.0001$ for all datasets, $n$ = 5-6). For minor and major workers of *C. pennsylvanicus*, dorsal/ventral surface had a significant effect on sensillum counts (two-way ANOVA, $p=0.0034$ for minors and $p<0.0001$ for majors). Bonferroni's multiple comparisons test was used for post-hoc comparisons between dorsal/ventral abundance at each segment and $p<0.05$ is denoted by an asterisk.
